# Robust Online Multiband Drift Estimation in Electrophysiology Data

**DOI:** 10.1101/2022.12.04.519043

**Authors:** Charlie Windolf, Angelique C. Paulk, Yoav Kfir, Eric Trautmann, Samuel Garcia, Domokos Meszéna, William Muñoz, Richard Hardstone, Irene Caprara, Mohsen Jamali, Julien Boussard, Ziv M. Williams, Sydney S. Cash, Liam Paninski, Erdem Varol

## Abstract

High-density electrophysiology probes have opened new possibilities for systems neuroscience in human and non-human animals, but probe motion (or drift) while recording poses a challenge for downstream analyses, particularly in human recordings. Here, we improve on the state of the art for tracking this drift with an algorithm termed **DREDge** (**D**ecentralized **R**egistration of **E**lectrophysiology **D**ata) with four major contributions. First, we extend previous decentralized methods to exploit *multiband* information, leveraging the local field potential (LFP), in addition to spikes detected from the action potentials (AP). Second, we show that the LFP-based approach enables registration at *sub-second* temporal resolution. Third, we introduce an efficient *online* motion tracking algorithm, allowing the method to scale up to longer and higher spatial resolution recordings, which could facilitate real-time applications. Finally, we improve the *robustness* of the approach by accounting for the nonstationarities that occur in real data and by automating parameter selection. Together, these advances enable fully automated scalable registration of challenging datasets from both humans and mice.

## 1. INTRODUCTION

Dense electrophysiology via multi-channel microelectrode probes, such as the Neuropixels probe [1, 2], provide an unprecedented view of neural circuits in human and non-human animal brains at extremely high resolution both temporally (30 kHz) and spatially (20-400 μm). In contrast with older recording technologies using lower spatial resolution arrays, the latest probes that penetrate into the brain allow us to measure activity in large populations of neurons (several hundreds) and the local field potential (LFP) with high fidelity. Since their introduction and ongoing development, these probes have allowed testing a variety of novel scientific hypotheses, including those related to neural correlates of consciousness [3], motor planning [4] and visual choice tasks [5], cementing their role as a staple tool for systems neuroscience for the foreseeable future. Furthermore, Neuropixels probes have recently been successfully employed for high-quality intra-operative recordings in awake and anesthetized humans [6, 7], enabling us to directly answer fundamental questions about human brain physiology with possible clinical implications.

However, several biological and physical sources of noise and variability reduce the neural recording effectiveness of Neuropixels probes [8]. In the case of in vivo measurements, especially in human participants, the probe signal can be impacted by brain motion effects due to the heart rate and breathing of the patients as well as unexpected brain shifting during recording (such as when the participants start to talk in clinically indicated awake tasks) [6]. This motion results in drift in the voltage measurements across channels, potentially corrupting the ability to isolate single unit activity on a given channel, which may lead to undersampling of spikes or over-splitting of identified unit clusters [9, 10, 11]. Together, these errors reduce the ability to characterize the functional activity of neural populations that are measured.

While drift affects voltage measurements in any rigid electrode, it is both visible and fixable in probes with very high spatial resolution such as Neuropixels. There are two main approaches to solving the motion drift problem in dense electrophysiology probes. Experimental approaches involve designing hardware to steady the probe during measurement. For example, probe movement can be stabilized at the open craniotomy as done in non-human primate preparations using O-rings or other materials pressing down on the brain [12]. However, in the human operating room, attempts to stabilize the probe using the same techniques could induce problematic capillary damage on the surface of the cortex and may require considerable careful non-human testing before implementing these approaches.

Two main computational methods have been developed to estimate motion drift in Neuropixels probes. Kilosort 2.5 [2] uses a template-based approach, computing a template signal, and then using cross-correlations to shift the drifting signal back to the template space. In contrast, Varol et al. [13] take a *decentralized* approach, measuring local signal shifts between all pairs of time-binned signals to learn a global displacement vector. These two techniques have been in wide deployment by several experimental groups [6, 14], however, they have three main short-comings. Kilosort 2.5’s template-based approach is plagued by model misspecification if the neural signal rapidly changes, eliminating the possibility that the recording would be described by a single template. The decentralized approach [13] circumvents this issue but is hindered by the computation of a *T* × *T* matrix that might be prohibitively large in chronic recordings. Furthermore, both of these methods have a variety of parameters such as time bin sizes and correlation cutoff parameters that need to be carefully tuned by practitioners to recover the tracked motion, reducing their robustness in high-throughput settings. Last and most importantly, these two techniques rely on the spike waveforms, probe locations, and discrete spike times of the high-frequency action potential (AP) band of the voltage signal, but ignore the smoother local field potential (LFP) band to estimate drift. Although AP band approaches require localizing spikes over time to estimate motion, the number of spikes in smaller time bins becomes too sparse to accurately compute motion-induced shifts. Hence, AP band approaches are limited to estimate drift on the order of ~1 second temporal resolution. On the other hand, the LFP band possesses smoother and more continuous signal across the entire recording, which has the potential to capture drift with a temporal resolution that is only limited by the sampling rate of measurement (~2.5KHz).

To overcome these obstacles, we introduce a novel extension to the decentralized registration approach [13] with the following **main contributions**: 1) fast online GPU-based optimization of displacement estimates in large-scale recordings, 2) sub-second temporal resolution, 3) automatic statistical tuning of parameters, and 4) multiband (LFP and AP based) motion estimation. Together, these four new features provide us with a fast, scalable, and robust approach for motion-correcting electrophysiological data with minimal or no parameter tuning. We term our approach **DREDge** which stands for **D**ecentralized **R**egistration of **E**lectrophysiology **D**ata.

We validate **DREDge** on two human Neuropixels recordings [6] and a mouse recording [2] from two separate research groups. We measure its performance in terms of registration quality and run-time and compare it with state-of-the-art techniques [13] and [2]. See https://github.com/evarol/DREDge for open-source code.

## 2. METHODS

We first motivate the **DREDge** algorithm using the decentralized registration approach [13] that it builds on in section 2.1. Then we describe several additions that enable **DREDge** to estimate drift in multiband electrophysiology data in a robust, efficient, and online manner in sections 2.2, 2.4, and 2.5. In section 2.7, we describe the procedure for realigning electrophysiology data through interpolation, adjusting for the shifting signal after motion estimation. Finally, we summarize all of these steps in algorithmic pseudocode in section 2.6.

### 2.1. Review: decentralized registration

Given a *D* × *T* signal **R** with columns **r**_*t*_, our goal is to discover **p** ∈ ℝ^*T*^ such that **p**_*t*_ is the displacement of the tth time bin **r**_*t*_. Here, **R** may be the signal from the LFP band, or it may be a rasterized representation of spiking activity from the AP band 1. In the LFP case, in practice, rather than using LFP directly, we spatially filter it to obtain one form of what we call current source density (CSD) [15], which provides spatially sharpened local features for registration. In the AP case, we construct **R** by first estimating depth positions along the probe for all spikes, typically using a point-source localization model [16]. Then, we divide the depth domain into *D* bins and the time domain into *T* bins and set **R**_*dt*_ to the average amplitude of spikes falling in the dth depth and tth time bins, taking the average in empty bins to be 0 by convention. In the LFP case, we construct **R** by directly taking the voltage signal at time *t* at each channel location with depth *d* such that **R**_*dt*_ is the LFP value at depth *d* and time t (see Fig.1 for examples of **R** in the left side).

**Fig. 1.**
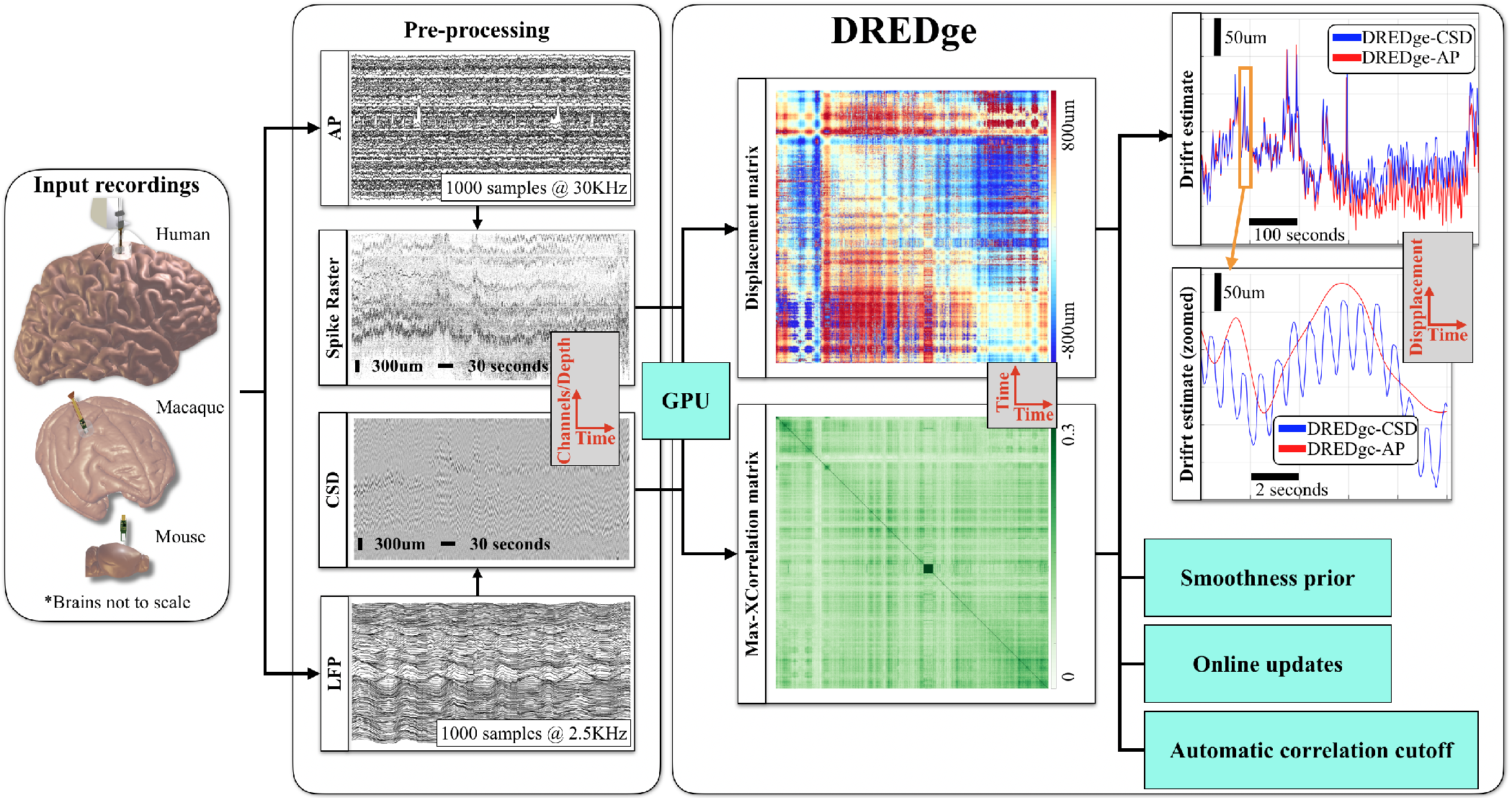
Overview of the **DREDge** algorithm. AP or LFP data from electrophysiology recordings from human, non-human primate, or mouse data is first processed to yield depth (D) × time (T) features **R**. Then each time-binned feature is cross-correlated with every other time-binned feature to generate a *T* × *T* displacement shift matrix and its corresponding maximum crosscorrelation. This step is done efficiently on the GPU. The displacement matrix is filtered using an automatically derived correlation cutoff and the remaining terms are used to solve a *centralization* equation to estimate drift estimates for each time bin. This procedure is robustified using priors that ensure that nearby drift terms are close to each other. Furthermore, for large recordings, the entire routine is done in smaller overlapping time chunks in an online fashion to efficiently calculate drift without the need to store a large *T* × *T* displacement shift matrix in memory.

The decentralized approach [13] infers **p** using estimates of the displacements between all pairs of time bins, represented in a *T* × *T* antisymmetric matrix **D** with

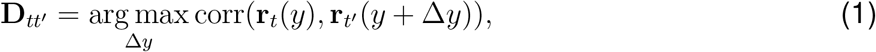

the spatial offset which maximizes the correlation between the two time bins. For AP-band problems, *T* is measured in seconds and this matrix is relatively small; for longer recordings or LF bands, the online method below avoids large matrices. The “centralization” problem, then, is to find

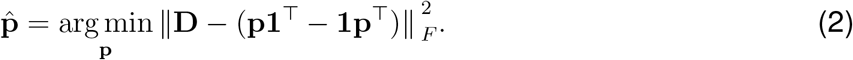

The solution to this simple version of the problem is the row mean:

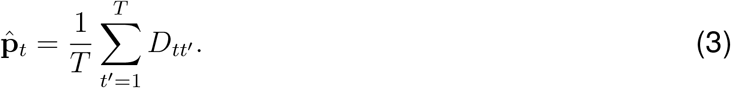

### 2.2. Extension: decentralized registration with correlation-based subsampling

If due to nonstationarities in the neural activity or large amounts of drift, two time bins **r**_*t*_ and **r**_*t′*_, do not contain the same features, then their pairwise displacement **D**_*tt′*_, should be excluded from the objective in Eq. (2). For example, this could occur in two time bins that are temporally distant, and the subject has completely different neural subpopulations firing, preventing a good shift to be found that overlays these two patterns.

Thus a simple heuristic approach is to include only those pairs of time bins whose maximal normalized cross-correlation exceeds a certain threshold, which indicates similar neural activity patterns up to a shift. To that end, we fix a correlation threshold *θ*, and let **C**_*tt*′_, = corr(**r**_*t*_(*y*), **r**_*t*′_, (*y* + *D_tt′_,*)), to be the correlation corresponding to the displacement estimate **D**_*tt*′_. Let **S** ∈ ℝ^*T*×*T*^ be the thresholded correlation matrix with 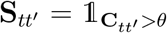. Now, we modify Eq. (2) to its subsampled form:

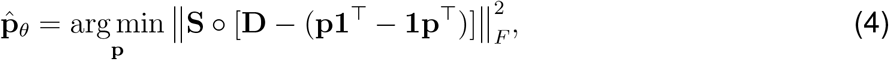

where ○ indicates the elementwise product.

To efficiently solve this problem, consider the set of pairs of times 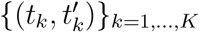, where *K* = ∑_*t,t′*_, *S_tt′_*, such that 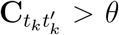. Now, let 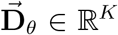 be the vector whose *k*th element is 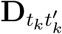, and let the matrix **A** ∈ ℝ^*K*×*T*^ have elements 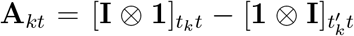. Then, Eq. (4) can be rewritten in the least squares form

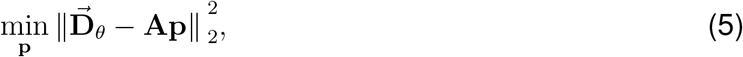

which can be solved by a sparse least squares solver (e.g. LSMR [17]). Note that such least squares systems usually require *O*(*T*^3^) to solve, but since **A** only has 2*K* << *T*^2^ non-zero elements, the complexity is reduced to *O*(*K*^2^*T*).

### 2.3. Adaptive choice of subsampling threshold

Since different modalities and recordings will have different statistics, it is important to find a way to set the correlation threshold robustly. One simple solution is to set this threshold to a low percentile of the distributions of maximum cross-correlations between neighboring time bins. Since almost all neighboring time bins contain the same features, this method will discover a correlation threshold that is suitable for non-neighboring time bins in a way that adapts to the characteristics of the recording. In the online method below, a threshold can be chosen in this manner for each new batch in order to adapt to nonstationarities in the signal.

### 2.4. Extension: smoothing prior for robustness

Since some time bins may be poorly correlated with the others, these bins may be separated from or only weakly linked to the rest through the subsampled cost function. This effect can lead to jumps in the resulting 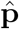. To resolve this issue, we place a standard Brownian prior on 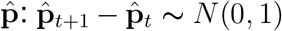. Since the difference operator is linear and sparse, it is simple to incorporate this prior into the above least squares framework, effectively turning our objective function into:

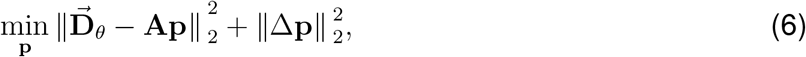

where Δ is the *T* × *T* finite difference matrix with elements Δ_*t,t*_ = 2 and Δ_*t,t*–1_ = Δ_*t*-1,*t*_ = −1.

### 2.5. Extension: online motion tracking

The methods in the previous two sections scale at least quadratically in *T*, since we compute *T* × *T* matrices **C** and **D** and solve a *T* × *T* system. When registering spiking data, where the time bins typically have lengths on the order of 1 second, this is no issue except in very long chronic recordings, or when registering sub-second resolution drift in human patients using the CSD or LFP signal.

To mitigate this, we estimate drift in chunks in an ‘online’ fashion. First, break the data **R** into *C* chunks of size at most *D* × *T*_0_, **R**^(*c*)^, *c* = 1,…, *C*. We initialize the algorithm by using the batch version of **DREDge**(Algorithm 1) to find **p**^(1)^, the displacement in the first block. Then, given the previous chunk’s displacement estimate **p**^(*c*)^, we can find the current chunk’s displacement estimate **p**^(*c*+1)^ according to the problems in Eqs. (2) and (4). Proceeding through the recording chunk by chunk, we can recover the full displacement estimate by concatenating those in each chunk. Since the sizes of the chunks’ sub-problems are bounded, this method will scale *linearly* in the total length of the recording. We present the online method in the case without correlation-based subsampling, since the notation for the subsampling will complicate things unnecessarily, but the extension is direct as in Sec. 2.2 and our results use subsampling.

Let 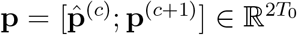 be the concatenation of the two displacement vectors, and define these chunks’ 2*T*_0_ × 2*T*_0_ displacement matrix

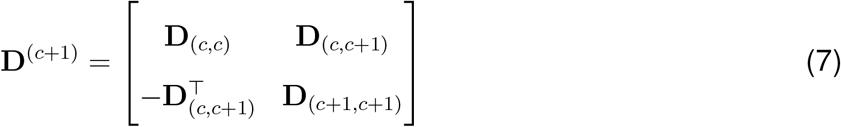

where the *T*_0_ × *T*_0_ blocks **D**_(*c,c*)_, **D**_(*c*;*c*+1)_, and **D**_(*c*+1,*c*+1)_ are pairwise displacement estimates in the previous block **R**^(*c*)^, between the two blocks **R**^(*c*)^ and **R**^(*c*+1)^, and within the current block **R**^(*c*+1)^, respectively. Consider Eq. (2), modified to hold 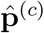 fixed:

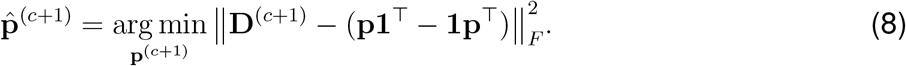

This online registration problem ensures that 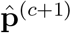 both aligns with the past estimate and centralizes the pairwise displacement estimates for the current time bins. Removing terms which do not include **p**^(*c*+1)^, Eq. (8) simplifies to

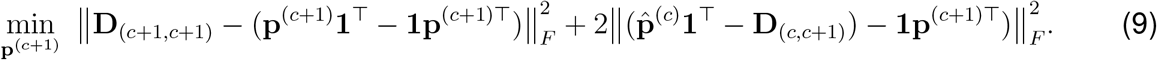

Here, the first term is the usual decentralized objective, ensuring the fidelity of the estimate in the current block. The cross term pushes **p**^(*c*+1)^ towards the column means of 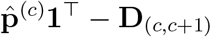, encouraging alignment with the past. As above, we can incorporate subsampling and solve this problem by rewriting it in OLS form and using a sparse least squares solver.

### 2.6. Algorithmic details

We summarize the above pipeline to estimate motion drift in electrophysiology data as the **DREDge** algorithm whose pseudocode details can be found in Algorithm 1. Note that the spatiotemporal signal matrix **R** ∈ ℝ^*D*×*T*^ can either be based on the AP band or the LFP band, resulting in two versions of the algorithm that we refer to below as **DREDge**-AP and **DREDge**-CSD. Algorithm 1 denotes the “batch” version of motion estimation. In large data cases such as in chronic recordings or in real-time applications, Algorithm 1 can serve as a *subroutine* for an online estimation of motion as described in section 2.5 where smaller time chunks of signal matrices **R** act as input in a streaming fashion.

**Algorithm 1.**
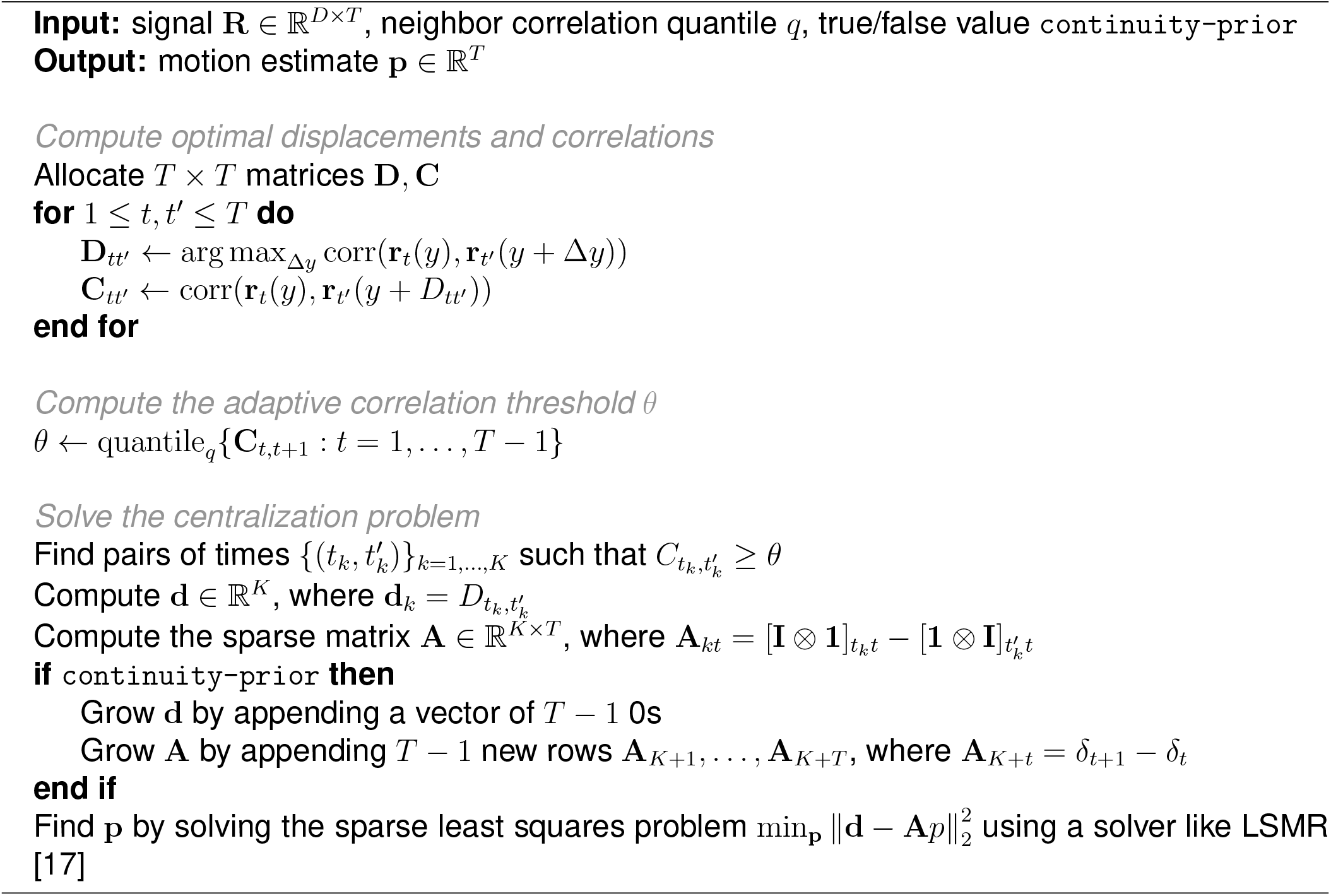
DREDge (batch).

### 2.7. Data interpolation after motion estimation

After estimating the motion trace using DREDge, this displacement estimate can be used to apply motion correction to the underlying raw data using a variety of interpolation strategies.

For example, the open source SpikeInterface [11] library can be used to interpolate the underlying raw data to correct for the motion, as in Fig. 4. SpikeInterface makes several strategies for interpolation available, including nearest-neighbor interpolation, kriging, and an inverse-distance weighted interpolation strategy. We used the latter, which we will outline here: first, a displaced coordinate is computed for each channel at each time according to the drift estimate, leading to a set of time-varying coordinates for a virtual probe. If the displacement at a given time was zero, then the virtual probe is not offset, so the original recording is used at that time. Otherwise, each virtual channel’s value is set to a weighted average of the three channels closest to its displaced location, where the weights are the inverse distances from the virtual channel to the three channels’ unregistered positions.

In Fig. 5, a similar alternative approach was used which can be more convenient for pipelines built around KiloSort and Matlab. This approach extends KiloSort 2.5’s kriging interpolation [2], enabling it to correct motion estimates sampled at arbitrary frequencies, rather than being locked to KiloSort’s temporal block length. Source code for this approach is available at https://github.com/williamunoz/DREDge_Interpolation.

## 3. RESULTS

### 3.1. Motion correction metrics

Experiments were carried out in three datasets: a mouse dataset with induced motion (dataset1 from [2]), and two human datasets (Pts. 02 and 09 from [6]). The mouse dataset is characterized by a slow (period ~ 100s) and shallow (tens of microns) induced triangle wave drift, while the human datasets are characterized by large (hundreds of microns) and fast (sub-second) drift driven by heartbeat and breathing patterns. Comparisons were carried out against the registration algorithm in Kilosort 2.5, which performs well on the same mouse dataset as previously shown in [2], but fails on the human datasets (see Fig. 2.(a, c)). Online methods used 10^4^ samples per chunk.

**Fig. 2.**
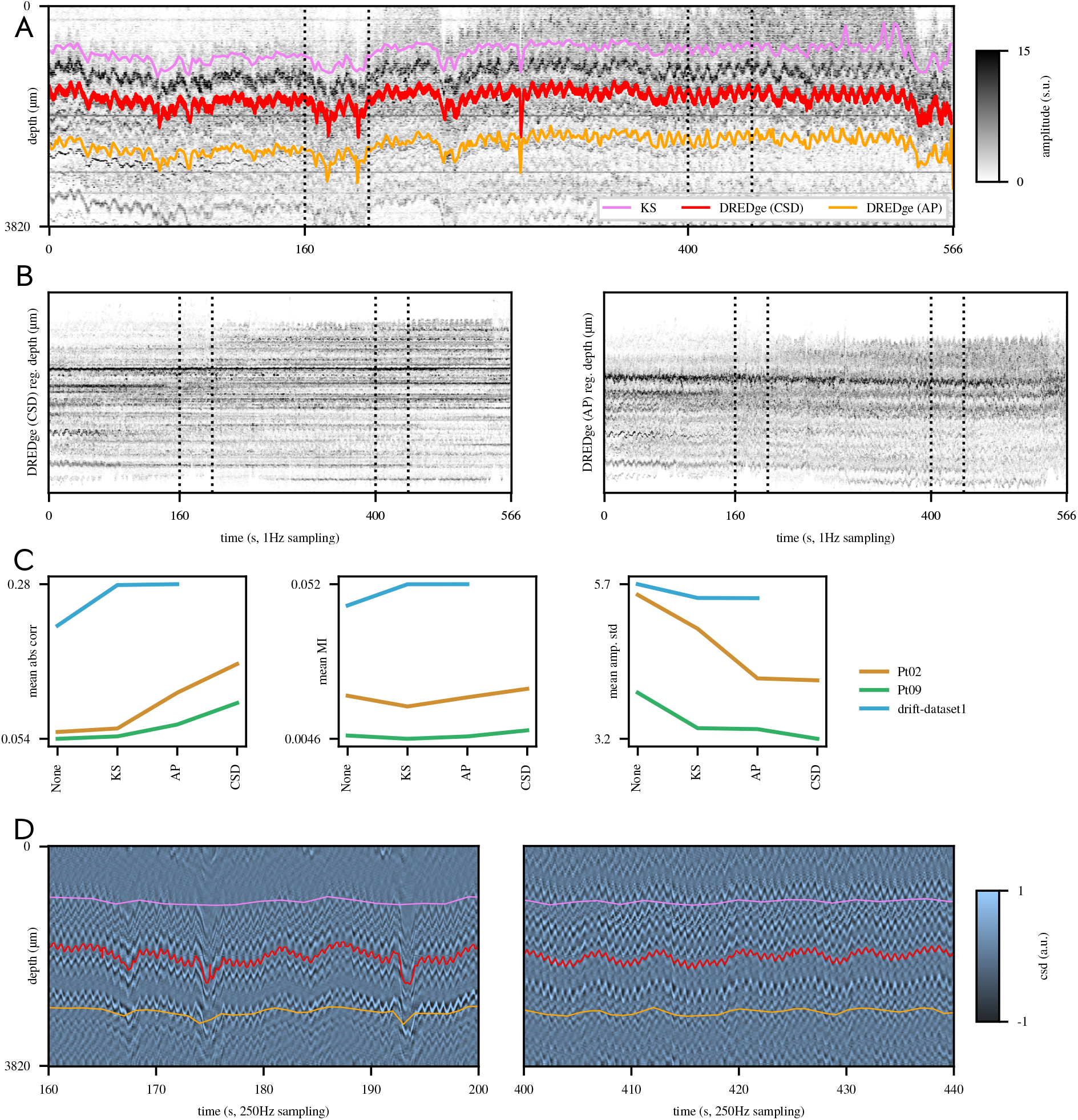
(a) Rasterized spike amplitudes from the Pt. 02 dataset [6], with estimated motion traces from Kilosort 2.5 and **DREDge**; regions of interest starting at 160 and 400s displayed in (d) show that the large jumps do correspond to the observed spatially filtered LFP which we refer to as CSD (d). (b) Spike amplitude maps after shifting the spike depths according to **DREDge** displacement estimates (CSD, top, and AP, below). (c) Metrics on three datasets, two human (Pts. 02 and 09, [6]) and one mouse (drift-dataset1, [2]). The first two metrics are the mean correlation/mutual information across pairs of rasterized spike activity time bins; the last is the mean standard dev. of the amplitude in-depth bins. (d) Motion traces for 40s regions of interest in the CSD. Color scale shared in (a) and (b).

Due to the characteristic noise present in unsorted spiking data, the AP methods (KS and **DREDge**-AP) shown in Fig. 2 cannot be pushed to a sub-second resolution and thus cannot capture the fine drift discovered by the **DREDge**-CSD method (Fig. 2.c), reflected in a well-stabilized spike amplitude raster (Fig. 2.a, top) when compared to the AP method (Fig. 2.a, bottom). The CSD method’s improvements over the AP method are reflected in its performance on simple metrics (Fig. 2.c).

### 3.2. Computational cost and parameter sensitivity

While the time complexity of the AP-based method is on par with Kilosort and both are just a few seconds (Fig. 3.a, top), when running on CSD sampled at 250Hz the runtime increases dramatically, an effect which is mitigated by the online method (Fig. 3.a, bottom). By using the prior and automatic correlation thresholding, the CSD method can robustly register full recordings (Fig. 3.d), without incurring shrinkage from the prior (right). Intuition for the effect of the prior is examined in a simulation study (Fig. 3.b), in which randomly selected time bins in the mouse AP data are zeroed out, resulting in “glitches” in the correlation and displacement matrices (right). With no prior, these glitches carry through to the drift estimate, but the prior is able to remove them. Throughout these figures, adaptive correlation thresholds (Section 2.3) were used. The 5th percentile of correlations of neighboring frames is used in AP, where the time scale is faster and thus more nonstationarities appear, while the 0.1 percentile is used in the smoother CSD. These thresholds are among the best performing when looking at a grid of choices (Fig. 3.c).

**Fig. 3.**
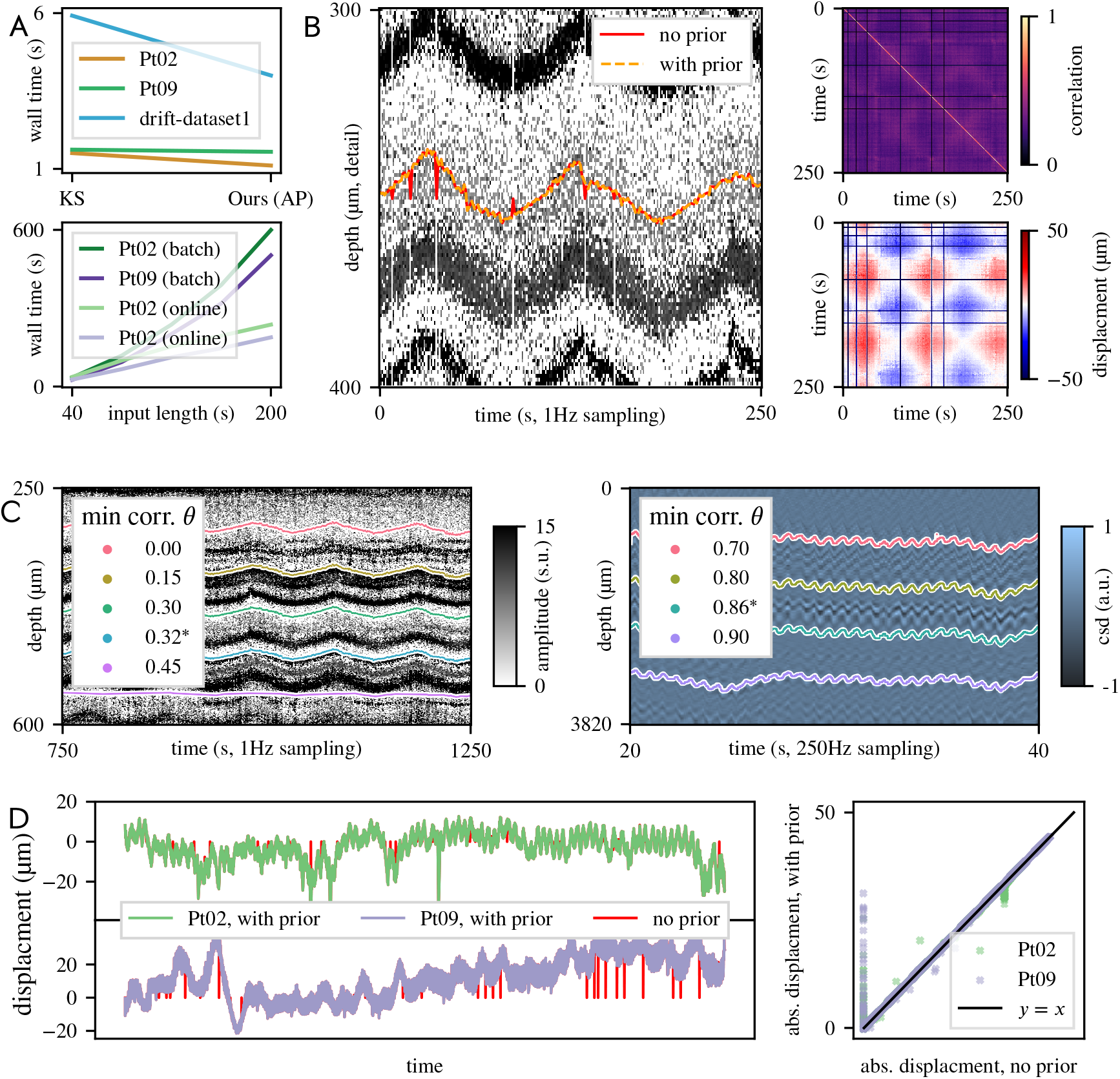
(a) Timing comparisons: top, **DREDge**-AP method vs. KS; below, online vs. offline in CSD. **DREDge**-AP is on par with KS thanks to a fast GPU implementation. (b) Comparing results with no prior and with prior on a test version of drift-dataset1 [2] with 5% of time bins erased at random: left, estimated motion traces over rasterized spike amplitudes; right, correlation and displacement matrices; horizontal and vertical stripes in these reflect the erased time bins. (c) Motion estimates using adaptive correlation threshold choices (marked *) vs. a grid of other choices: top, in real mouse AP data (*θ*\* = 5th percentile of neighbor correlations) and bottom, in human CSD (0.1 percentile). (d) Left, full CSD displacement estimates with prior (purple, green) and without (red); right, adding the prior pushes estimates for isolated frames away from 0 without causing shrinkage.

### 3.3. Validating DREDge by motion correction of localizations

To further validate our drift estimate, we shifted localization features [16] of detected spikes according to the drift estimated at the time of each spike event. Clusters that were not visible at first (Fig. 4.a) appear after correction (Fig. 4.b). In particular, spikes from a particular cluster shown in green in panel (b) have localizations with a broad spread before motion correction. To see if these particular spikes truly correspond to a single unit, we interpolated the binary file to correct for the drift using the open-source SpikeInterface framework [11]. As expected, waveforms corresponding to spikes in the cluster were spread across many channels in the uncorrected recording (Fig. 4.d), but appear as a stable unit after correction (Fig. 4.e), validating that the drift correction was accurate. For another perspective, we check to see if the cluster can be identified in the localizations of spikes detected in the interpolated binary. Indeed, after localizing events from that binary, shown in (Fig. 4.c), we identified the same cluster manually (purple dots). Waveforms loaded at times corresponding to these spikes (Fig. 4.f) match those shown in (Fig. 4.e). Further, after using SpikeInterface [11] to identify matching spike times across these clusters, the Venn diagram in (Fig. 4.g) shows that these clusters have high spike time agreement.

**Fig. 4.**
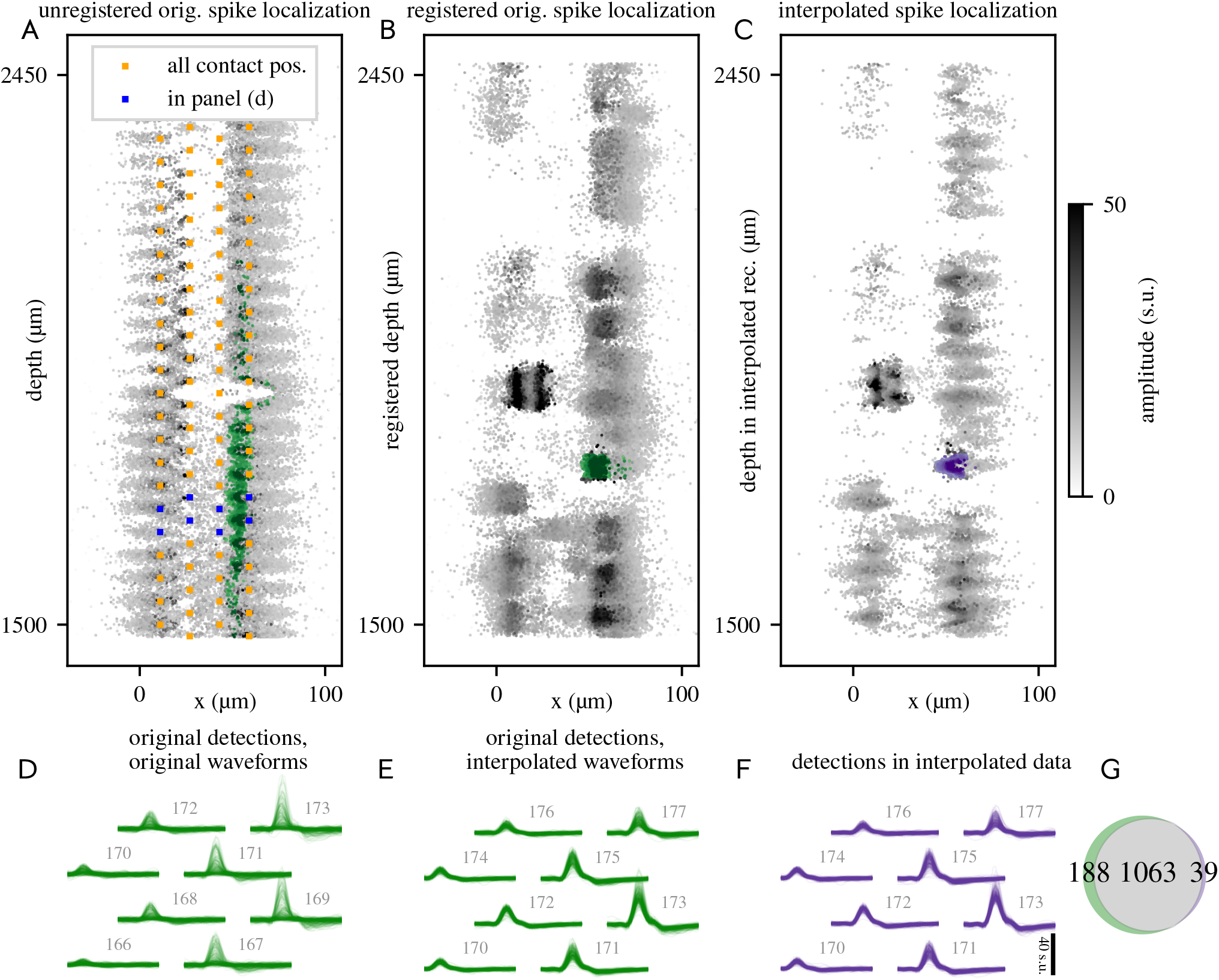
Checking the drift estimate using drift-corrected spike localizations [16] and interpolated waveforms. (a) Localizations of spikes detected in the Pt. 02 dataset. (b) Spike localizations shifted according to a drift estimate (**DREDge**-CSD in Fig. 2.a); green dots in (a) and (b) represent the same events, corresponding to spikes in a well-isolated cluster in (b) isolated by manually thresholding amplitude and selecting a rectangular region in *x, z*. (c) Spike localizations computed from events detected in a version of the recording which was interpolated to correct for drift using the SpikeInterface framework [11]. Purple dots lie in a region matching the region used to select the green dots in (b). (d) Waveforms corresponding to spikes highlighted in the green box in (a-b) read from the original recording, shown on high-amplitude channels (contacts shown in blue in (a)). (e) Waveforms corresponding to the same spike times, read from the motion-corrected interpolated binary. The cluster which emerged after correcting the spike depths corresponds to a unit with a well-stereotyped waveform shape in the interpolated binary, providing a validation of the drift estimate. (f) Waveforms corresponding to spikes highlighted in purple in (c); these waveforms match (e) in appearance. (g) A Venn diagram comparing spike times found in the green and purple clusters in (b) and (c); most (1063) spikes are shared between the two spike trains, and few spikes in each cluster do not match (188 and 39 spikes).

**Fig. 5.**
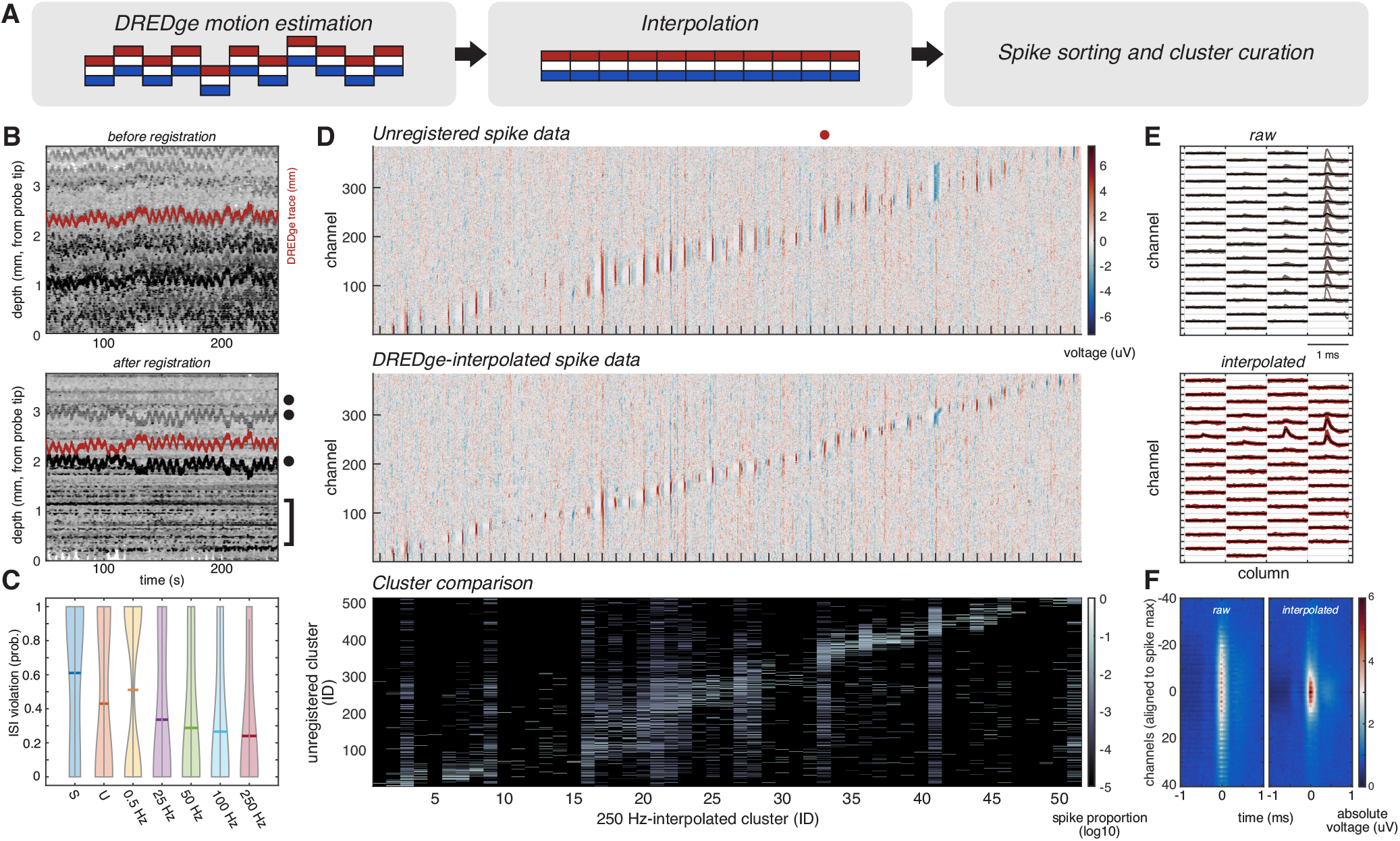
Interpolation with DREDge motion estimate of human intraoperative Neuropixels spiking data. (a) Schematic of preprocessing steps for spiking data. (b) Spike detections (normalized amplitude gray scale) before (top) and after (bottom) registration with an optimized DREDge motion estimate (red). Note the emergence of spike bands on bottom panel (bracket), as well as the propagation of DREDge anti-correlated single channel noise (dots). (c) Progressive decrease in inter-spike interval violation probability with increasing interpolation rate (0.5 - 250 Hz), as compared to unregistered data (U) and interpolation with a scrambled DREDge trace (S). Bar represents mean. (d) Full probe spatial distribution of spikes in non-interpolated condition (top) and 250 Hz-interpolated clusters (middle). (Bottom) Comparison of 250 Hz interpolated spike assignments to unregistered clusters, showing over-splitting and cross-contamination of unregistered clusters. (e) Representative unregistered (top) and 250 Hz-interpolated (bottom) unit (red dot on panel D). Scale bar 1 ms. (f) Average spatial distribution of spike clusters when non-interpolated (left) and 250 Hz-interpolated (right).

### 3.4. Data interpolation after DREDge motion estimation

To optimize DREDge motion correction in human multichannel recording with Neuropixels probes, we interpolated the estimated DREDge motion (Fig. 5.a) and observed progressively improved alignment with faster interpolation rates (0.5 to 250 Hz; Fig. 5.b). As motion artifacts in human intraoperative recordings have multiple frequency components, we optimized the DREDge interpolation rate to be well above the Nyquist limit for respiration- (0.25-0.3 Hz) and heartbeat-related (1-1.7 Hz) motions (Fig. 5.b). In these conditions, we observed the emergence of aligned spike bands that resemble well-isolated units (Fig. 5.b, bracket). Furthermore, we also observed a spatially restricted propagation of single-channel noise (Fig. 5.b, dots), following the DREDge trace, which was easily identifiable during cluster curation.

We then employed KiloSort 1 for spike sorting of unregistered and 250 Hz-interpolated data. This choice of clustering algorithm corresponded to our intention to apply a clustering method with no inherent motion correction to better isolate and evaluate the outcome of our DREDge motion correction on spiking data. In agreement with the observed spike bands after interpolation, we concomitantly observed that increasing interpolation rates progressively decreased inter-spike interval violation probabilities (Fig. 5.c). Of note, using a too-low interpolation rate of 0.5 Hz yielded an inter-spike interval violation probability distribution similar to unregistered data, emphasizing the importance of correcting a broad band of motion frequencies.

Furthermore, when we compared the spatial profile of spikes in DREDge-interpolated clusters before and after alignment, we observed an expected spatial tightening of spike distributions after alignment (Fig. 5.d-e). The spatial extent of our aligned spike clusters is consistent with prior Neuropixels data [18], demonstrating that average spike spatial distribution of cortical neurons spans an average of 0.2 mm (Fig. 5.f).

We then compared the cluster assignments from unregistered and 250 Hz-interpolated data (5.d, bottom). We observed that the spikes assigned to virtually every realigned cluster corresponded to spikes found on multiple unregistered clusters, suggesting that the unregistered spike data is over-split by clustering algorithms. Conversely, we noted that virtually every unregistered cluster was made up of spikes from multiple registered clusters, importantly suggesting a significant degree of contamination of unregistered clusters, consistent with their higher inter-spike interval violation probabilities. These over-splitting and contamination effects further emphasize the importance of motion correction in this dataset for the isolation of single units.

## 4. DISCUSSION

Here, we present an online decentralized algorithm, **DREDge**, to track drift in electrophysiological recordings from the brain using Neuropixels probes. We demonstrate the applicability of tracking the movement of the neural signal relative to the probe with this approach in three different data sets, two recordings in the human cortex [6] and one in mouse [2]. **DREDge** is shown to work using as input both action potentials, representing the activities of single neurons, as well as the local field potential, representing the summed activity of hundreds to thousands of neurons. We develop methods to adapt this algorithm to the structure of individual datasets, improving the robustness and ease of use of this tool.

Finally, we suggest that **DREDge** could be used to improve the clustering of spike localization features for applications in spike sorting. Furthermore, as a fast online estimation of the motion of the probe relative to the neural signal, **DREDge** has potential applications in recording sessions with real-time neural feedback and brain-computer interfaces (BCI) [19, 20].

The latter point may be a key feature in gathering neural data using Neuropixels in the human cortex. Unlike in mouse preparations [1, 2], recordings in humans, so far, have presented a challenge in that the brain, by virtue of the scale and size, moves considerably due to the heart-beat, breathing, the participants talking, etc. This has required researchers in this domain to rely on manual tracking or only use data where the spike rate is high enough to track the neural signal reliably using Kilosort tools [6, 7]. Such other approaches might not fully correct for the observed motion, with implications for lower yields of unit quality and/or number. **DREDge** allows researchers to move forward in using such precious data to understand human brain activity.

A major question is whether after applying the tracked motion to the signal, there are changes that are not present in the original signal. Our data suggest that applying **DREDge** correction to motion-degraded spike and local field potential data has the potential to accurately reconstruct these electrophysiological signals, while not altering their spatiotemporal profiles in datasets without significant motion artefacts. However, the power of our approach is that we use both the LFP and the spiking activity in parallel to estimate the motion and, alternately, can use each, particularly the spike waveforms, to confirm that we are not adding in artefacts. Further validation could therefore make use of 2D cross-correlation to determine if new spiking waveforms were found in the corrected data sets [6]. An additional question is whether ongoing large-scale neural changes such as evoked potentials due to a task, clinically indicated neural signals such as epileptiform activity, anesthesia-induced burst suppression [6], or other types of stimulation could alter the motion registration and, therefore, the tracking. Further analyses of additional data sets from humans, non-human primates, rodents, and other species where we know specific events are occurring will be necessary to parse this effect in a future study.

Finally, a true test of this approach is how it can handle different types of probes, probe designs, and layouts, both Neuropixels and other silicon probes, and how this can alter spike sorting [1, 2, 21]. To this end, we were able to apply our method to the staggered Neuropixels 1.0 version (the two human data sets) and the Neuropixels 2.0 probe (the mouse data set). Future work would involve applying the method to other probe layouts, particularly to ask whether this approach could be applied to low spatial resolution probes and the spatial and temporal limitations in tracking motion relative to the neural signal.

To facilitate this sort of portability, **DREDge** is being developed into a part of the SpikeInterface [11] framework. In future applications, and as **DREDge** is applied to further data sets using different probe layouts and reaching deeper brain structures other than the cortex, we propose that the underlying algorithm may be applicable to a number of different conditions and paradigms within and beyond electrophysiology.

## 5. ACKNOWLEDGMENTS

We would like to thank Yangling Chou, Aaron Tripp, Brian Coughlin, and Fausto Minidio for their help in the original data collection. We would like to especially thank the patients for their willingness to participate in this research. EV is supported by 1K99MH128772-01A1. This research was supported by the ECOR and K24-NS088568 (to SSC) and the Tiny Blue Dot Foundation (to SSC and ACP), NIH grant U01NS121616 (to ZMW), and NIH/NINDS Neuroscience Resident Research Program R25NS065743 (to WM). CW, EV and LP are funded by Simons Foundation 543023, NSF Neuronex Award DBI-1707398 and the Gatsby Charitable Foundation. The views and conclusions contained in this document are those of the authors and do not represent the official policies, either expressed or implied, of the funding sources. The funders had no role in the study design, data collection and analysis, decision to publish, or preparation of the manuscript.

## 6. COMPETING INTERESTS STATEMENT

The MGH Translational Research Center has clinical research support agreements with Neuralink, Paradromics, and Synchron, for which SSC provide consultative input. None of these entities listed are involved with this research or the Neuropixels device. The remaining authors declare no competing interests.

## Notes

https://github.com/evarol/dredge

https://github.com/SpikeInterface/spikeinterface

### Summary of Updates

Added a new study of the effect of DREDge on human intraoperative Neuropixels spike sorting results using interpolation to correct for the motion (WM), shown in the new Figure 5. Methods, Results, and Discussion are updated to incorporate this new analysis.

